# SeqScreen-Nano: a computational platform for rapid, in-field characterization of previously unseen pathogens

**DOI:** 10.1101/2023.02.10.528096

**Authors:** Advait Balaji, Yunxi Liu, Michael G. Nute, Bingbing Hu, Anthony Kappell, Danielle S. LeSassier, Gene D. Godbold, Krista L. Ternus, Todd J. Treangen

**Affiliations:** Department of Computer Science, Rice University, 6100 Main Street, Houston, TX, USA; Signature Science LLC, 8329 North Mopac Expressway, Austin, TX, USA; Signature Science LLC, 1670 Discovery Drive, Charlottesville, VA, USA

## Abstract

The COVID-19 pandemic forever underscored the need for biosurveillance platforms capable of rapid detection of previously unseen pathogens. Oxford Nanopore Technology (ONT) couples long-read sequencing with in-field capability, opening the door to real-time, in-field biosurveillance. Though a promising technology, streaming assignment of accurate functional and taxonomic labels with nanopore reads remains challenging given: (i) individual reads can span multiple genes, (ii) individual reads may contain truncated genes, and pseudogenes, (iii) the error rate of the ONT platform that may introduce frameshifts and missense errors, and (iv) the computational costs of read-by-read analysis may exceed that of in-field computational equipment. Altogether, these challenges highlight a need for novel computational approaches. To this end, we describe SeqSeqscreen-Nano, a novel and portable computational platform for the characterization of novel pathogens. Based on results from simulated and synthetic microbial communities, SeqScreen-Nano can identify Open Reading Frames (ORFs) across the length of raw ONT reads and then use the predicted ORFs for accurate functional characterization and taxonomic classification. SeqScreen-Nano can run efficiently in a memory-constrained environment (less than 32GB of RAM), allowing it to be utilized in resource-limited settings. SeqScreen-Nano can also process reads directly from the ONT MinlON sequencing device, enabling rapid, in-field characterization of previously unseen pathogens. SeqScreen-Nano (v4.0) is available on GitLab at: https://gitlab.com/treangenlab/seqscreen

## INTRODUCTION

Long-read sequencing technology has come of age, and was recently deemed the 2022 method of the year. Indeed, it is rapidly becoming the preferred choice for genomic and metagenomic analyses over short-read sequencing technologies^1,2^. In contrast to shorter sequence fragments, longer reads offer improved resolution and accuracy across several tasks such as de novo assembly^3,4^, structural variant identification^5,6^, metagenomic assembly^7^, and microbiome analyses^8,9^. The availability of portable single-molecule sequencers such as the Oxford Nanopore MinION and SmidgION have enabled real-time in-field sequencing of biological samples, drastically reducing the time for analysis^10^. These handheld sequencers can connect to a laptop or a smartphone for downstream analyses, enabling powerful in-field sequencing and analysis potential. However, rapid and accurate processing of nanopore sequencing in a memory-efficient and resource-limited setting remains a challenge^11^. Nonetheless, in-field nanopore sequencing analyses have proven essential for time-sensitive applications such as infectious disease surveillance. They have been used to track the spread Zika^12^, Ebola^13^ and COVID- 19^14^ outbreaks. These applications of in-field nanopore sequencing provide a foundation for improved in-field biosurveillance and tracking of previously unobserved pathogens^15^. Previously unseen pathogen detection from in-field isolates remains under-explored^16^. Recently, several novel approaches for real-time analysis, including MAIRA^17^, CRuMPIT^18^,and npReader from Japsa package^19^ have been developed. However, some challenges still exist for in-field utilization of these tools with MinION sequencers. MAIRA is a GUI-based tool to analyze batches of nanopore reads out a sequencer for sample composition. While MAIRA is optimized for genus level taxonomic profiling, it can output species level profile on-demand. Functional information from MAIRA is limited to functional marker genes obtained from RefSeq. MAIRA also reports CARD^20^ and VFDB^21^ annotations, however, we have previously shown that VFDB annotations only capture limited mechanisms of pathogenicity^22^. While npReader is more efficient at handling batches of high throughput nanopore data, it includes no information on pathogenic function and, while important, is limited to reporting on antibiotic resistance. Finally, while CRuMPIT can provide taxonomic profiles of in-field samples from nanopore sequencing data and is based upon Centrifuge^23^ it lacks functional information.

Most approaches to identifying pathogens in a metagenomic sample rely on taxonomic classification and similarity to reference databases. MetaMaps^9^ and MEGAN-LR^24^ are two of the most widely used tools for metagenomic characterization using long reads. MetaMaps is a long-read taxonomic classifier and composition estimator that performs the classification in two stages. In the first step, MetaMaps uses MashMap to align reads to a modified RefSeq database. In the second step, classification is carried out by running the initial mappings through several iterations of an Expectation-Maximization algorithm. The algorithm re-assigns reads to the most probable taxonomic label based on alignment and mapping qualities. MetaMaps outputs both read-level and sample-wide taxonomic assignments, and provides a sample-wide compositional estimate. MEGAN-LR is a protein-based classification method that uses DIAMOND^25^ or LAST to map reads to the non-redundant protein database. The mappings are then processed by the meganizer module and taxonomy is assigned using the interval union Least Common Ancestor (LCA) algorithm which is a modified form of the canonical LCA algorithm that accounts for different taxonomic assignments for hits (or intervals) across the length of the read. MEGAN-LR also offers read-level functional assignments in various categories including InterPro^26^, eggNOG^27^ and KEGG^28^. However, accurate functional annotation of long reads is still challenging as no current tool exists for ORF identification^29^. Also, while these tools have achieved state-of-the art performance for taxonomic classification on long-read metagenomic datasets they are not suitable for characterizing novel emerging pathogens because of a lack of pathogen-specific functional labels. Further, their memory and time requirements of make them prohibitive for rapid in-field characterization of long reads in a streaming and resource-limited setting.

In our previous work, we presented SeqScreen, a novel pathogen screening platform that screened for pathogenic markers in individual reads obtained from any NGS sequencing experiment. In this prior contribution, we outlined a novel functional framework called Functions of Sequences of Concern (FunSoCs) that reports 32 different categories of pathogenic function and highlighted how they could be used for improving pathogen detection compared to standard taxonomic classifiers. In this work, we describe SeqScreen-Nano, which extends the FunSoC framework to long reads and can be used to identify pathogen genomes within metagenomes. Specifically, we present three main contributions; First, using simulated nanopore reads we show that SeqScreen-Nano can sensitively identify ORFs in long reads leading to accurate functional characterization at read-level compared to closely related tools like MMseqs2^30^and MEGAN-LR. Second, we present a novel algorithmic approach based on the minimum-set cover and reference-based inference to retrieve the correct species present in the sample. Third, using appropriate parameters and efficient data structures SeqScreen-Nano is more computationally and memory efficient than MetaMaps and MEGAN-LR. This enables SeqScreen-Nano to run on laptops and directly process data from portable nanopore sequencers resulting in shorter turn-around times for real-time analyses. We show that the FunSoC framework and the reference inference approach can help alleviate challenges pertaining to low-abundance pathogens. Additionally, we highlight the advantage of the FunSoC framework over taxonomic classifiers in identifying novel emerging pathogens. Table 1 outlines the contributions made by SeqScreen in context to previous tools.

**Table 1:** Microbial community profiling and taxonomic classification tools designed for long reads. Taxonomy = represents the targeted taxonomic classification level, Function - Functional assignments and gene ontology information, Novel Pathogen - Identification of pathogenic markers capable of detecting novel pathogens, and Streaming - Can process batches of reads from ONT sequencer in an “online”fashion. *GUI based tool. ^*a*^ Optimized for Genus but can report Species (on demand). ^b^Optimized for Species but reports Strain.

| Tool | Taxonomy | Function | Novel Pathogen | Streaming |
| --- | --- | --- | --- | --- |
| Centrifuge<br>(Kim et al. 2016) | Species/Strain | No | No | No |
| CRuMPIT<br>(Sanderson et al. 2018) | Species/Strain | No | No | Yes |
| MetaMaps<br>(Dilthey et al. 2019) | Species/Strain | No | No | No |
| MEGAN-LR<br>(Huson et al. 2018) | Genus/Species | NCBI nr | No | No |
| MAIRA*<br>(Albrecht et al. 2020) | Genus/Species<sup>a</sup> | RefSeq marker proteins | VFDB + CARD | Yes |
| **SeqScreen-Nano**<br>**(This work)** | **Species/Strain<sup>b</sup>** | **UniProt** | **FunSoCs** | **Yes** |

## METHODS

### Pipeline overview

The SeqScreen-Nano pipeline is based on the SeqScreen pipeline with substantial additions to deal with the complexity of long-read sequences Briefly, the SeqScreen pipeline is built upon five different workflows; (i) Initialize, (ii) SeqMapper, (iii) Protein and Taxonomic Identification, (iv) Functional Annotation and, (v) SeqScreen report generation. Workflows (ii) and (iii) are exclusive to the *sensitive* mode of SeqScreen while the other workflows are shared with the *default* mode. For more information on the different modes and workflows within SeqScreen we point the readers to^22^. SeqScreen-Nano leverages the SeqScreen FunSoC framework and features several enhancements, which we will now detail.

First, we develop a reference-guided ORF prediction step for long reads and contigs. We accomplish this by using DIAMOND^25^ to align the input to the custom UniRef100 databases (proteins with annotation in a two-step mapping process and it is run using the parameters -*range-culling*, -*frameshift 15 and* -*max-target-seqs 1*. This enables DIAMOND to return multiple hits along the length of the entire read rather than hits from a specific region of the read. We call these hits *candidate ORFs*. The ORFs are tracked and extracted efficiently using an interval tree data structure. The *–min-orf* parameter controlling the minimum length of hit is set to 30. The regions between the candidate ORFs with length ≥ 90 base pairs that were unmapped during the first mapping step are then mapped to the second DIAMOND database which contains proteins with lower annotation scores.

Second, a new step called the *reference inference* module is used to call the presence or absence of a particular taxon in the sample. The reference inference module can be run in two user-controlled modes; *online* by using the *–online* flag or *offline* by default. Putative taxonomic ids from the SeqScreen taxonomic identification stage are used to obtain reference genomes of the potential species in the sample. In online mode, current reference genomes are downloaded from NCBI; whereas, in offline mode, the custom BSAT database associated with the SeqMapper workflow is used as the source of possible reference sequences. Once the reference sequences for the species are obtained, all the reads are mapped using Minimap2 to each of the individual reference genomes (independently) and the depth and breadth of coverage are calculated. In the subsequent step, high-coverage reference genomes are concatenated and the reads are remapped to the concatenated reference to simulate “read stealing” where reads have a chance to map to the best reference in the presence of similar references (i.e., closely related species or strains). The coverage scores are recalculated after the second mapping step and are used to call species presence.Third, we have added a streaming capability capable of processing reads as they are base called off of an ONT MinION sequencer, capable of producing pathogen calls within user-defined increments in time (2 hours, 4 hours, etc) and within resource-constrained computational environments. Additional details on the streaming mode are detailed below.

### Read level taxonomic characterization

We present a novel algorithm for read-level taxonomic classification in long reads based completely on ORF annotations. We first try and select the most probable taxonomic assignment using a simple majority voting scheme. If most assignments are at the genus level and there exists at least one assignment to a species of one of the genera, then the respective genus is counted as a species hit. In case a simple majority does not exist, then a weighted minimum set cover algorithm is used to assign taxonomy id to the read. The minimum set cover algorithm is formulated as follows. Given the set of all UniRef100 hits in the sample given by Equation 1 where each *u_i_* represents a UniRef100 hit (for each ORF) and *u_s_p__* refers to individual subsets pf UniRef100 hits or the set of UniRef100 hits for each read in Equation 2.

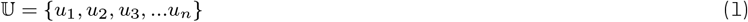

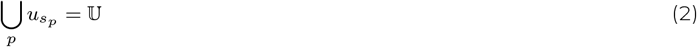

We can then formulate a reverse mapping of taxonomic ids to UniRef as given in Equation 3 and define cost c_i_ for each taxonomy *t_i_* in Equation 4.

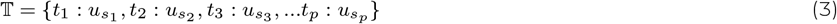

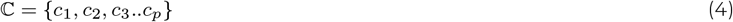

The goal is then to find a subset 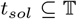 that covers 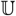 such that the total cost is minimized 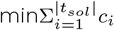. Reads with ambiguous hits are assigned the taxonomic id of the ORF present in *t_sol_* and if multiple exist, then the one with the lowest e-value is chosen.

### Calculation of coverage score and Average Nucleotide Identity

The depth of coverage is calculated by using *samtools depth* and the observed breadth of coverage (O) is determined by calculating the number of times each position is covered by reads at the given depth and is normalized by the genome length. We then calculate the expected breadth of coverage using a probabilistic estimate as follows. We first define *M* as the mean mapping length in Equation 5, which is the total depth at each position normalized by the number of reads mapped (denoted by X).

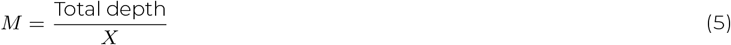

Further, we partition the genome into buckets of size equal to the mean mapping length and call it *N*. We can then calculate the expected value of mapping (E) and the standard deviation (stdev) by Equation 6 and Equation 7, respectively.

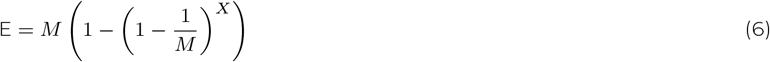

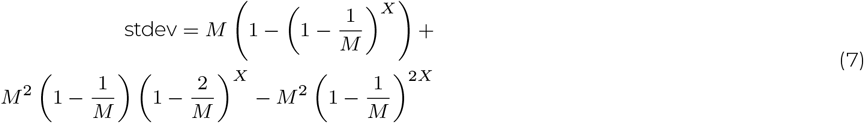

The expected value of mapping is then normalized by M to get the expected coverage. The coverage score (CS) is then defined as the ratio of the observed breadth of coverage and expected breadth of coverage as given by Equation 8.

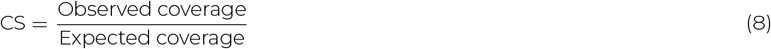

The Average Nucleotide Identity (ANI) is calculated using the *samtools consensus* output and is the ratio of the number of bases matched to the total number of positions. If the ANI is greater than 0.97 then the species is determined to be present. Species is also marked as present if the change in coverage score is ≥ −0.1 and ANI is greater than 0.9, but if the coverage score is between 0.7 and 0.9 then the genus is reported as present (i.e., not enough confidence for species call). If none of these conditions are satisfied, then the species is marked absent. The reference inference pipeline and rubric used to call the presence of species and genera is illustrated in Figure 1.

**Figure 1:**
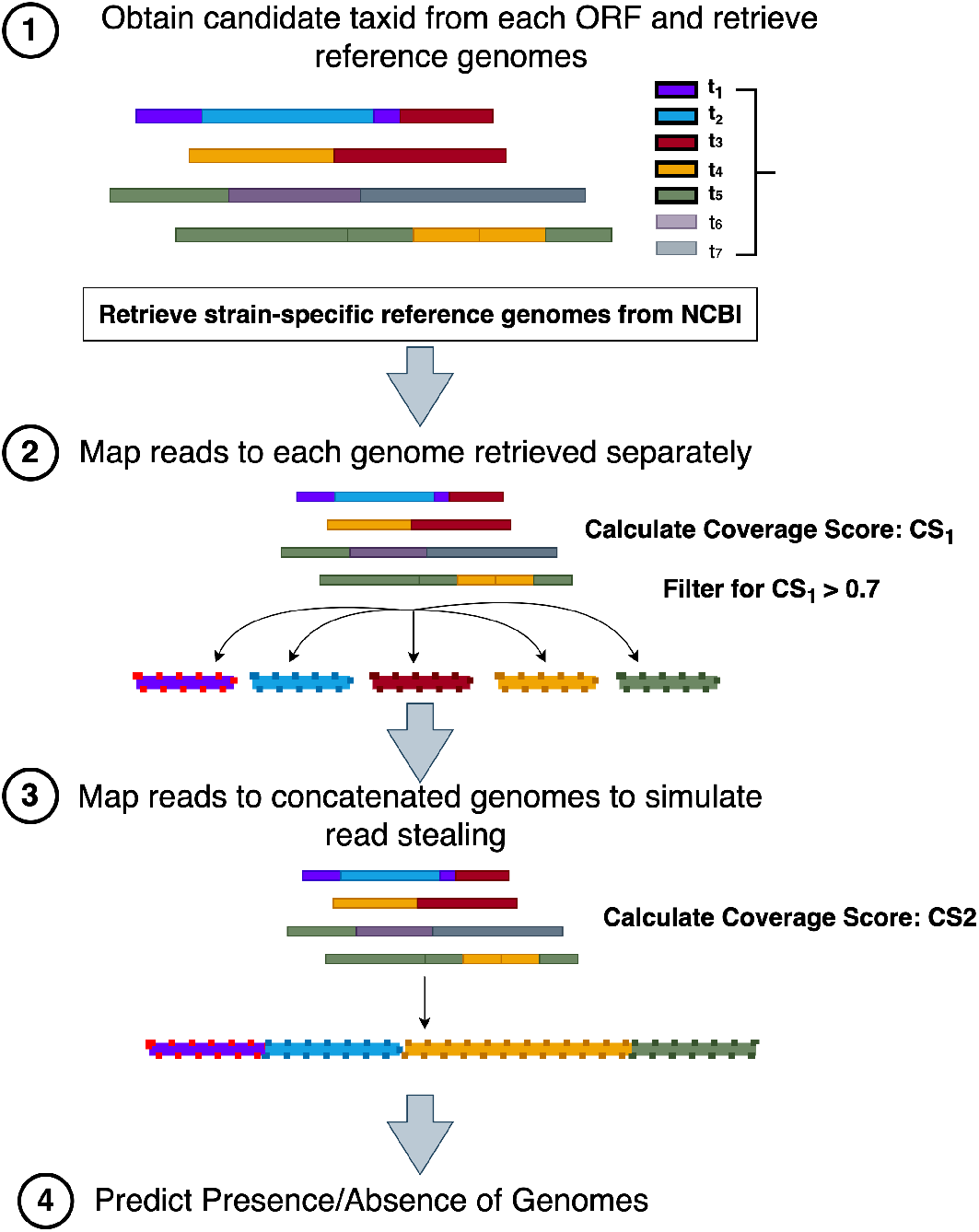
Reference Inference pipeline. Workflow of mapping and prediction stages using reference inference for calling sample-wide species or genus present.

### Streaming mode in SeqScreen-Nano

A novel feature of SeqScreen-Nano is the ability to process batches of reads output by the sequencer (e.g., MinION) in a streaming setting. This is achieved by having a wrapper script that calls SeqScreen-Nano internally. During a MinION run, the wrapper periodically checks for new FASTQ files being generated in the output directory. The directory is scanned every 60 seconds to check for new files. The new files are then added to a queue. By default, if the number of reads in the queue is 200000, then a SeqScreen-Nano run is triggered. If no new file is seen after 30 minutes, then the remaining reads in the queue are collected and processed, and the script exits.

### Datasets and Tools

#### Identification of Open Reading Frames (ORFs) in nanopore reads

To test the accuracy of ORF identification in long reads, we simulated reads from *Clostridium botulinum* and *Influenza A* (*H3N2*). For the former, the genome was split into 10 bins covering the length of the genome and 5 reads were simulated from each bin. For the latter, 5 reads were simulated from each segment. Reads were simulated using NanoSim-H^31,32^ with a minimum length of 1000bp and default error rate used in the *ecoli_R9_1D* profile. MMseqs2 and MEGAN-LR were chosen for comparison to SeqScreen-Nano. ORFs were predicted from SeqScreen-Nano and MMseqs2 based on hits in a read with unique query start and end pairs after alignment. ORF information for MEGAN-LR was obtained by analyzing the number of COGs obtained after alignment by DIAMOND and meganizer for long reads using MEGAN6.

#### Identification of pathogens present in metagenomes

We used two distinct mock metagenome communities to test sample-wide genus and species identification. The ZymoBIOMICS Microbial Community DNA Standard consisted of 10 distinct species, including 8 bacteria and 2 fungi, and was downloaded from SRA accession ERR3152364. The downloaded FASTQ file was subsampled randomly, resulting in a total of 174181 sampled reads representing all 10 species at varying abundance. Sequencing of the ZymoBIOMICS Gut Microbiome Standard was carried out internally in our laboratory. Sequencing library preparation was performed using the Oxford Nanopore Technologies Rapid Sequencing Kit (RAD004) as per the manufacturers instruction. DNA input was 400 ng for the ZymoBIOMICS Microbial Community DNA Standard (Zymo Research, D6306) and extracted DNA from the ZymoBIOMICS Gut Microbiome Standard (ZymoResearch, D6331). The ZymoBIOMICS Gut Microbiome Standard was extracted using ZymoBIOMICS DNA Miniprep Kit (ZymoResearch, D4300T). The ZymoBIOMICS Microbial Community DNA Standard II (ZymoResearch, D6311) was loaded with 165 ng DNA, the maximum input of 7.5 μL at 22 ng/ μL. The library was loaded onto a primed MinION Flow Cell, R9.4.1. The flow cell was housed in the MinION Mk1C. The settings of the sequencing run were set for default values for the sequencing kit and fast basecalling using the onboard guppy basecaller (6.1.5) with Q8 quality cutoff for pass and failed reads and outputting both fast5 and FASTQ files at 4000 reads per file. Modified settings included removing the reserving of pores (Off or 0%) and size setting to minimum 20 bp, as an extreme for testing that could allow for size exclusion in post run quality control. A total of 100000 reads from the first 25 batches of output from the MinION were considered for the experiment.

#### Identification of low-abundance pathogens in metagenomes

To test detection and identifications of low-abundance pathogens in metagenome, we spiked the subsampled ZymoBIOMICS Microbial Community DNA Standard with reads from *Clostridium botulinum* covering the breadth of the genome. The abundance of *Clostridium botulinum* in the sample was 0.08%, which was the least abundant of all the microbes present in the sample.

#### Identification of novel pathogens

Transfer of genomic segments was simulated in three pairs of near-neighbor species, and the reads were analyzed to compare SeqScreen-Nano with various taxonomic classifiers. The experiment aimed to evaluate if the included tools could identify a novel pathogen that evolved due to gene loss/gain events or mutations. Transfer of sequences was simulated by extracting 1000 segments from the more pathogenic near-neighbor chosen at random to be either 300bp, 500bp or 700bp in length. A total of 15000bp reads (minimum length 500bp) were simulated using NanoSim for the merged genome (after gene transfer) and compared to those simulated from less pathogenic species. The near-neighbor pairs used in this experiment were (i) *E.coli O157:H7* and *E.coli K12*(ii) *Staphylococcus aureus* and *Staphylococcus epidermidis* and (iii) *Bacillus anthracis* and *Bacillus cereus*. In (iii), instead of transferring segments of the genome between backbones, plasmids in *Bacillus cereus* were replaced by those from *Bacillus anthracis*.FunSoCs were predicted using SeqScreen-Nano and compared to taxonomic classification results obtained from MetaMaps and Centrifuge. The FunSoCs *antibiotic resistance* and *secretion* were excluded from the SeqScreen results to better highlight more pathogenic FunSoCs.

#### Analysis of Fecal Microbiota Transplant (FMT) sample

As a reasonable negative control of our FunSoC framework within SeqScreen-Nano, we set out to it on a screened and cleared FMT healthy Donor sample. To accomplish this, we used a FMT Donor sample we previously characterized^33^. Briefly, two pediatric patients with a recurrent *Clostridioides difficile* Infection (CDI) diagnosis received FMT under IRB-approved informed consent (#H-31066) at Baylor College of Medicine. Patients received filtered, frozen-thawed fecal preparations from a standardized donor (3840 year old male during donations) via colonoscopy. The donor screening and fecal preparation procedures were approved by the U.S. FDA (IND#15743). Shotgun metagenomic sequencing was performed with 200 ng of input DNA as previously described^34^ and was submitted to NCBI BioProject database PRJNA743023. The paired-end reads were then assembled into contigs using MEGAHIT with *–k-min 31* and *–k-max 91*. This resulted in contigs with an average length of 1396 bp and median length of 496 bp.

## RESULTS

### Identification of ORFs within long reads

Since long reads may contain multiple ORFs within a single read it is imperative for accurate functional characterization that tools can identify the correct number of ORFs per read. To test this, we simulated reads from *Influenza A (H3N2)* and *Clostridium botulinum*. For *Clostridium botulinum*, The length of the genome was partitioned into 10 bins and we simulated 5 reads from each bin. For *Influenza A (H3N2)*, we simulated 5 reads from each of the eight segments. To compare ORF assignments, we considered MMseqs2 and MEGAN-LR. It is worth noting here that both SeqScreen-Nano and MEGAN-LR use DIAMOND as a mapping step, but we have optimized parameters of DIAMOND whereas MEGAN-LR is run with default parameters as presented in the manual as well as by prior studies. In SeqScreen-Nano, we identify an ORF as having a unique (start,end) tuple after alignment and use the same metric for SeqScreen-Nano. For MEGAN-LR, we take the number of eggNOG annotations after meganization as the number of ORFs As seen in Figure 2 (A.) we observe that in both *Influenza A (H3N2)* as well as *Clostridium botulinum*, SeqScreen-Nano is closer to the actual number of ORFs per read as compared to MMseqs2 and MEGAN-LR. We see from both examples that MMseqs2 seems to overpredict ORFs, whereas MEGAN-LR underpredicts ORFs. Further Figure 2 (B.) describes the mean, median and standard deviation of difference between actual and observed ORF counts across all the reads.

**Figure 2:**
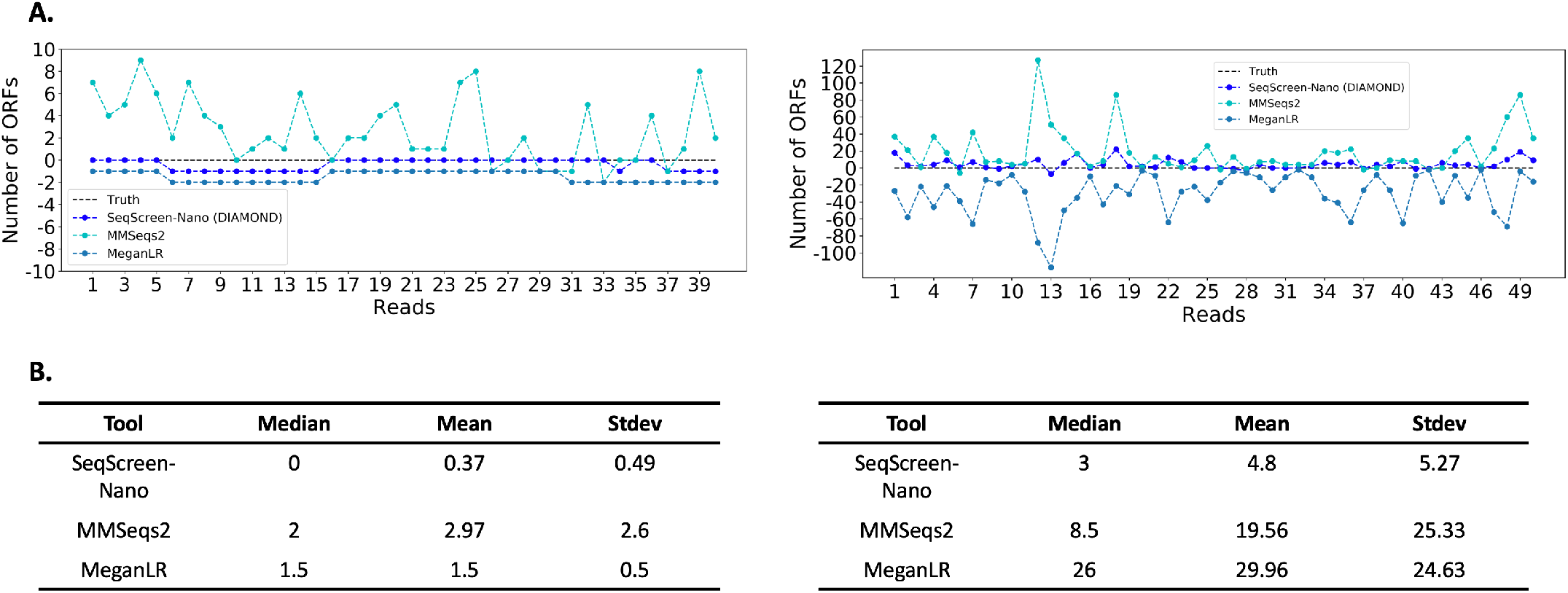
Comparison of reference guided ORF detection across nanopore reads for *Influenza A (H3N2)* (left) and *Clostridium botulinum* (right) (A.) Number of ORFs reported by each tool as compared to the truth. Panel B shows the mean, median, and standard deviation from the truth.

### Identification of pathogens in metagenomes

In addition to functional and pathogenic labels at the ORF level, it is relevant to report likely taxonomic labels present in the sample, especially if previously seen pathogens are present in the sample. To demonstrate SeqScreen-Nano’s capability to identify pathogens within metagenomes, we tested it on two subsampled mock metagenome communities - the ZymoBIOMICS Microbial Community DNA Standard (10 microbes, 10 species) and ZymoBIOMICS Gut Microbiome Standard (21 microbes, 17 species). Since, we were only interested in comparing at genus and species levels, we collapsed all the 5 different *E.coli* present in the ZymoBIOMICS Gut Microbiome Standard to the species level and counted it the same in our analysis. We compared SeqScreen-Nano to two other popular long-read taxonomic classifiers; MetaMaps and Centrifuge. Further, as both MetaMaps and Centrifuge calculate sample-wide abundances and read counts for specific taxonomic levels, unlike SeqScreen-Nano that directly reports *Presence/Absence*, we set a threshold of 0.001 for both MetaMaps and Centrifuge to call *Presence/Absence*. We can see from Figure 3 (A.) that on the ZymoBIOMICS Microbial Community DNA Standard, SeqScreen-Nano has perfect Precision (P) and Recall(R) at both genus and species levels, while Centrifuge has the worst performance of the three tools (P: 0.8, R:0.8 and P: 0.7, R, 0.7) respectively. MetaMaps fails to predict *Cryptococcus neoformans* at 0.001 abundance cutoff in the sample. In the ZymoBIOMICS Gut Microbiome Standard that we sequenced, we again observe from Figure 3 (B.) that SeqScreen-Nano has perfect precision at the Genus level compared to MetaMaps (0.86) and Centrifuge (0.71), whereas both SeqScreen and MetaMaps have similar recall at 0.76, followed by Centrifuge at 0.58. At the species level, all tools perform less accurately than at the genus level. However, SeqScreen-Nano still has the highest F1 score (0.66) compared to MetaMaps (0.57) and Centrifuge (0.43) due to its higher precision (0.76) at the species level.

**Figure 3:**
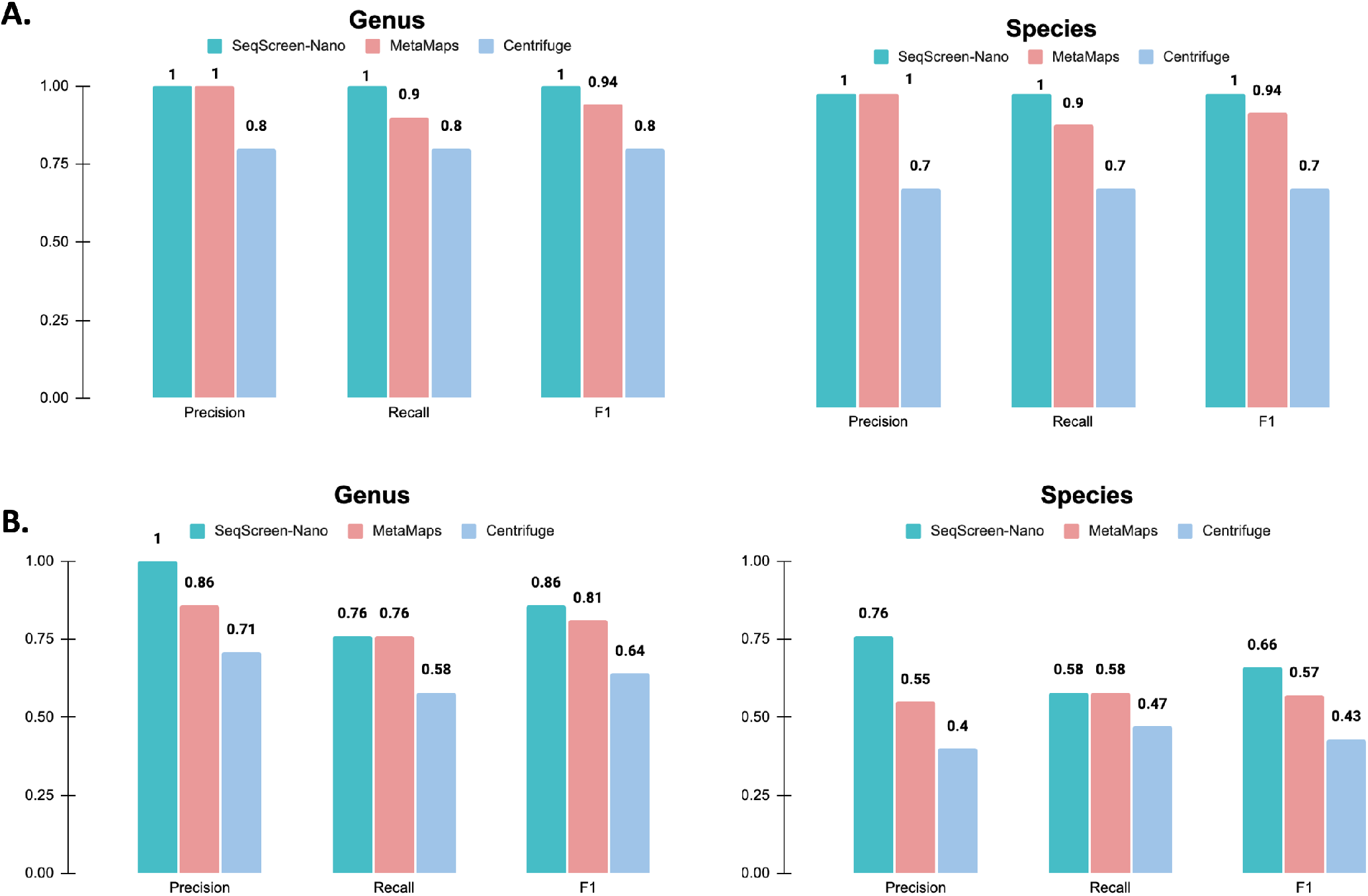
Taxonomic Performance on MinION sequences from ZymoBIOMICS Microbial Community DNA Standard and ZymoBIOMICS Gut Microbiome Standard. (A.) Precision, Recall, and F1 score at genus and species levels for ZymoBIOMICS Microbial Community DNA Standard. (B.) Precision, Recall, and F1 score at genus and species levels for ZymoBIOMICS Gut Microbiome Standard.

In addition to analyzing the accuracy of different approaches, we also analyzed the computational requirements of each of these tools. Given the move towards portable sequencing, it is imperative that methods can run efficiently in a resource limited setting. Table 2 indicates the time (in minutes) and memory (in GB) for both of the ZymoBIOMICS mock communities. SeqScreen-Nano is the most memory efficient tool and the second fastest, while having the best accuracy in terms of precision and recall across both datasets. It is worth noting here that MetaMaps has a flexible maximium memory option to enable it to run in low memory settings, however, we have seen doing so causes a linear increase in run time. Since the major bottleneck for processing long-read data from portable sequencers is memory, SeqScreen-Nano is the only tool capable of accurately calling species level hits efficiently.

**Table 2:** Performance on MinION sequences from ZymoBIOMICS Microbial Community DNA Standard and ZymoBIOMICS Gut Microbiome Standard

| Sample/Tool | Time (minutes) |  |  | Memory (GB) |  |  |
| --- | --- | --- | --- | --- | --- | --- |
|  | SeqScreen-Nano | MetaMaps | Centrifuge | SeqScreen-Nano | MetaMaps | Centrifuge |
| ZymoBIOMICS Microbial Community DNA Standard (10 species) | 60 | 167 | 3 | 23.6 | 273.86 | 42.43 |
| ZymoBIOMICS Gut Microbiome Standard (17 species) | 39 | 115 | 3 | 19.44 | 273.86 | 40.59 |

### Identification of low-abundance pathogens in metagenomes

To elucidate the advantage of the FunSoC framework over canonical taxonomic approaches, we considered the case of a low-abundance pathogen. Reads belonging to *Clostridium botulinum* were spiked into the ZymoBIOMICS Microbial Community DNA Standard with 10 species at 0.08% abundance and covering regions across its genome. Figure 4 (A) shows the FunSoCs identified in the sample that were specific to *Clostridium botulinum* and the number of ORFs as reported by SeqScreen-Nano. Notably, we found hits to *disable organ* and *degrade ecm* FunSoCs associated with the *Botulinum neurotoxin type A* gene. We also identified a hit to the *Thiol-activated cytolysin* gene which was associated with the *cytotoxicity* FunSoC. To contrast FunSoC based pathogen identification against canonical taxonomy approaches, we compared against MetaMaps and Centrifuge. Figure 4 (B) shows precision and recall of these tools at 0.1% and 0.01% abundance thresholds at species level. We observe that at a higher threshold (0.1%) both MetaMaps and Centrifuge have high species-level precision and recall and could identify a majority of higher abundance microbes. However, when the abundance threshold was lowered to 0.01% to identify *Clostridium botulinum*, there was a significant drop in precision from 1.0 to 0.39 for MetaMaps and 0.72 to 0.18 for Centrifuge. While both tools identified *Clostridium botulinum* at this lower threshold, the increased number of false positives made the identification of the low-abundance pathogens ambiguous, in contrast to the FunSoC framework that could identify specific pathogenic genes present in *Clostridium botulinum*. Additionally, the reference inference module in SeqScreen-Nano identified *Clostridium botulinum* as present at the species level, underscoring the importance of breadth of coverage analysis to detect pathogens accurately at low-abundance.

**Figure 4:**
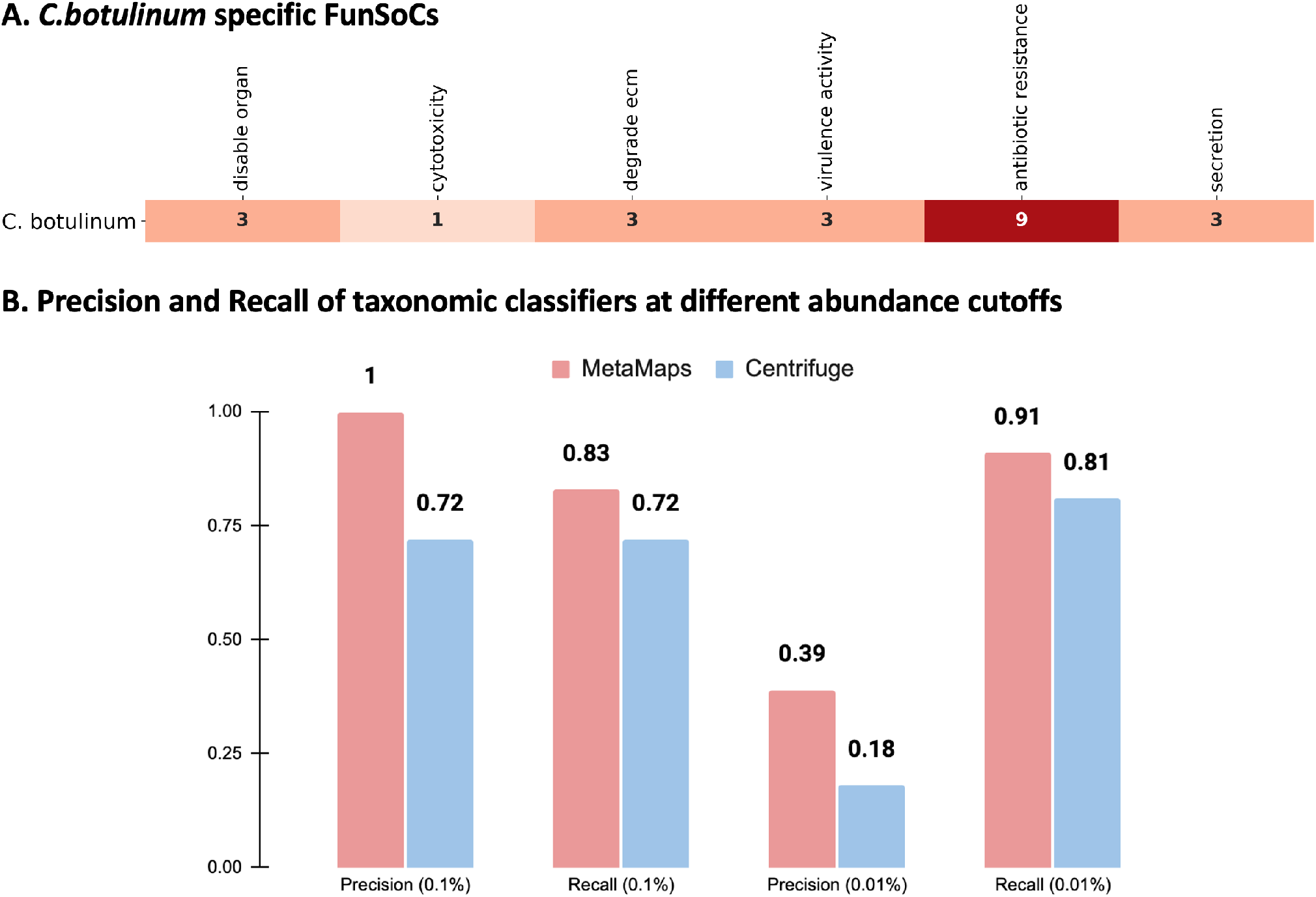
Performance on ZymoBIOMICS Microbial Community DNA Standard. (A.) FunSoCs reported by SeqScreen-Nano specific to *C.botulinum* in the spiked ZymoBIOMICS Microbial Community DNA Standard. Numbers represent the number of ORFs detected with that FunSoC. (B.) The Precision, Recall, and F1 score at genus and species levels for the spiked ZymoBIOMICS Microbial Community DNA Standard.

### Identification of simulated novel pathogens

We next consider a simulated experiment to describe the utility of FunSoCs in identifying novel pathogens over taxonomic approaches. Gene transfer events were simulated between three well-characterized pathogens and closely related species/strains namely (i) *E. coli* K12 and *E.coli* O157:H7, (ii) *Staphylococcus aureus* and *Staphylococcus epidermidis*, (iii) *Bacillus cereus* and *Bacillus anthracis*.

In cases (i) and (ii), random parts from the pathogen genome were merged into the commensal genome. In case (iii), plasmids from *Bacillus anthracis* were inserted in place of plasmids from *Bacillus cereus*. Figure 5(A) shows that when compared to unmutated *E.coli K12*, the merged *E. coli* genome had an increase in *cytotoxicity, secreted effector* and *virulence regulator* FunSoCs. The increase in *cytotoxicity* FunSoC was attributed to the introduction of *Shiga toxin* gene (*StxB*) from *E. coli* O157:H7. Additionally, we also observed a significant increase in the *secreted effector* FunSoC due to the presence of *secreted effector protein tir* gene. Both these genes are known to be absent in the *E. coli* K12 strain. Further, we found that increased copies of the *YhaJ* virulence regulator gene. In comparison, the top hits at the strain level for Metamaps and Centrifuge (threshold 0.005) were to *E. coli* K12 representing their most confident predictions, and only Centrifuge reported some presence of *E. coli O157:H7*, while most of its reads were predicted at the species level. (Figure 5(B) and Figure 5 (C)).

**Figure 5:**
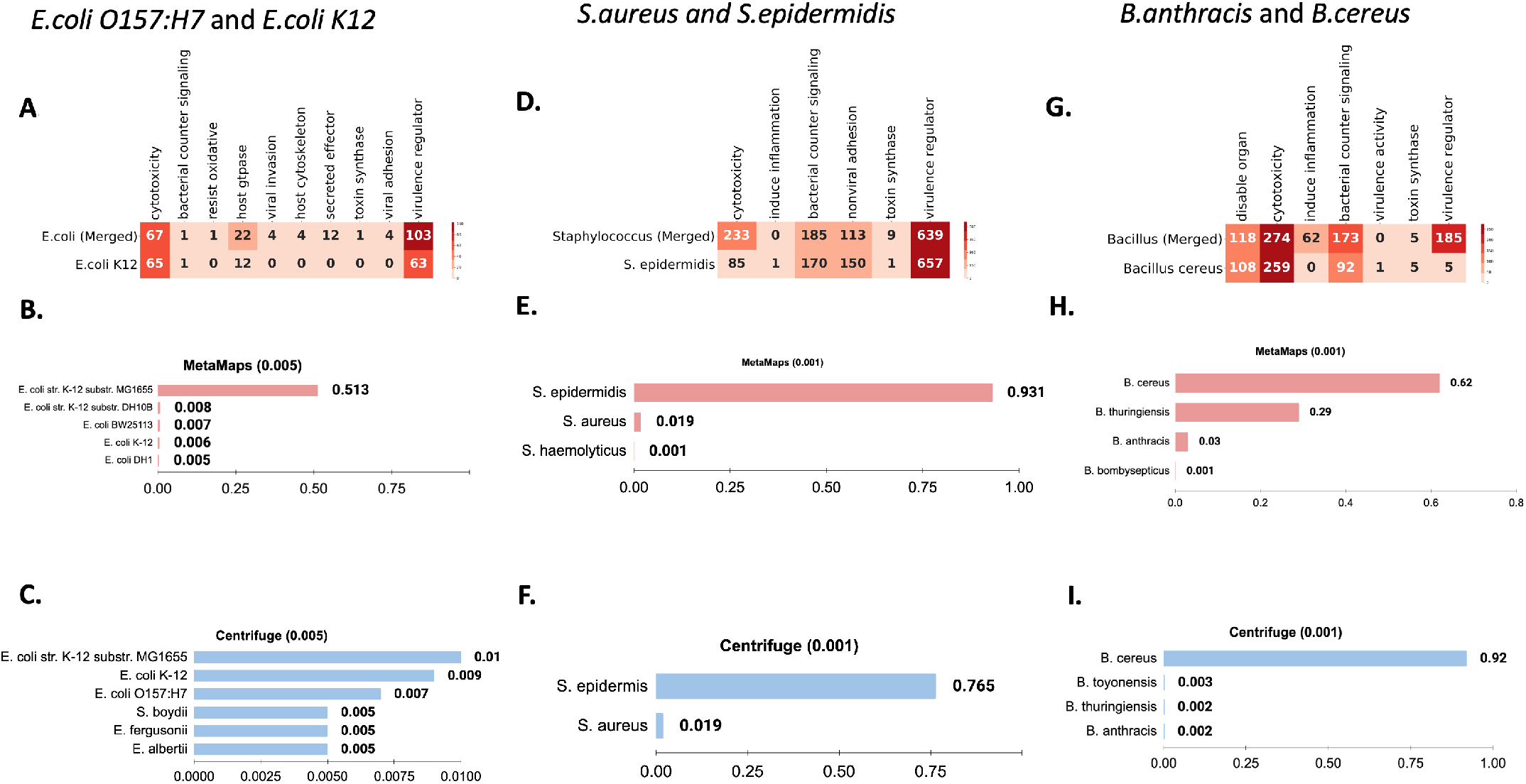
Analysis of simulated novel pathogens using SeqScreen-Nano, MetaMaps, and Centrifuge. (A.) FunSoCs for merged *E.coli* O157:H7 and *E.coli* K12 strains vs. *E. coli* K12. Numbers represent the number of ORFs detected with that FunSoC. (B.) MetaMaps results for merged *E. coli* genomes (C.) Centrifuge results for merged *E. coli* genomes (D.) FunSoCs for merged *S. aureus* and *S. epidermidis* vs. *S*. *epidermidis* (E.) MetaMaps results for merged *Staphylococcus* genomes (F.) Centrifuge results for merged *Staphylococcus* genomes (G.) FunSoCs for merged *B. anthracis* and *B. cereus* vs. *B. cereus* (H.) MetaMaps results for merged *Bacillus* genomes (I.) Centrifuge results for merged *Bacillus* genomes. The threshold for *E. coli* strains was set to 0.005 for better visualization.

In the case of *S.aureus* and *S. epidermidis*, we see an increased level of hits to the *cytotoxicity* FunSoC as a result of key *S.aureus* specific genes being identified, such as *Leucotoxin (LukE)* and *Gamma-hemolysin component C (HglC)*. Besides *cytotoxicity*, we also observed an increase in *bacterial counter signaling* and *toxin synthase* FunSoCs, with the latter being a result of additional hits to *Dihydrolipoyl dehydrogenase* and *Pyruvate decarboxylase* genes. In this case, we also observed increased hit in *S.epidermidis* compared to merged genome for the FunSoCs *nonviral adhesion, virulence regulator* and *induce inflammation* but the genes were found to ubiquitously present across different *Staphylococcus* species. As expected, both MetaMaps and Centrifuge both reported *S.epidermidis* as its top species level classification, followed by some trace presence of *S. aureus*.

Finally, we examined the case of replacing *B. cereus* plasmids with the highly pathogenic *B. anthracis* plasmids. We observed a slight increase in the FunSoCs *disable organ, cytotoxicity*, which contained *Hemolysin* and *Thiol-activated cytolysin* genes found in both species. In addition to these FunSoCs, there was a significant increase in *induce inflammation* and *virulence regulator*, which contained hits to genes like *Anthrax toxin expression trans-acting positive regulator, Capsule synthesis positive regulator (AcpB), Capsule synthesis positive regulator (AcpA)*, all of which were related to the pXO1 and pXO2 plasmids. SeqScreen-Nano was also able to identify the *Protective Antigen (PA)* gene in the merged genome which was appropriately associated with *bacterial counter signaling* and *induce inflammation* FunSoCs. In contrast, MetaMaps and Centrifuge predominately predicted *B. cereus* over *B. anthracis*, which shows that most taxonomic tools consider the higher proportion of reads from the backbone to make a species-level classification.

### Analysis of Fecal Microbiota Transplantation from Healthy Donor sample

As a final computational experiment, we evaluated SeqScreen-Nano’s performance on a real Fecal Microbiota Transplant sample. We used this sample both as a negative control (the expectation is that a screened and cleared FMT sample should contain few to no FunSoCs) and also to see if we could identify putative probiotic species within the sample. Specifically, we analyzed a healthy donor sample from BioProject PRJNA743023 that was cleared for FMT in patients with *C.difficile* infections. We analyzed the sample for probiotic bacteria as identified by the reference inference module. Specifically, among important genera that prevent recurrence of *C.difficile* infection we observed the presence of five species of *Blautia*, five species of *Roseburia*, two species of *Dorea* and one species of *Butyricicoccus* (positively identified for Genus). Previous studies have shown that these genera are known to be characteristic of super donors for FMT^35^. In addition to these, four species of the genus *Eubacterium* were also detected, which plays a key role in immunomodulation and suppression of inflammation^36^. At the species level, the donor sample contained *Akkermansia muciniphila* which is keystone probiotic with protective effects against *C.difficile* induced dysbiosis^37^. The FunSoC profile of the sample indicated extremely trace amounts of highly pathogenic FunSoCs and corroborated the profile of the donor sample.

## DISCUSSION

We described SeqScreen-Nano, a computational platform for rapid in-field characterization of previously unseen pathogens. SeqScreen-Nano builds off of our previously published SeqScreen tool in two key ways. First, it processes error-prone raw nanopore reads and leverages a reference-guided approach to identify multiple ORFs across the length of the read, as opposed to identifying a single best hit within low-error short reads. Second, in addition to performing detection of FunSoC-containing ORFs, SeqScreen-Nano performs reference inference of pathogens using our previously described algorithm, providing species-level and strain-level assignments. Our experimental results included a variety of simulated and synthetic datasets that highlighted encouraging SeqScreen-Nano taxonomic classification and pathogen characterization performance compared to state-of-the-art tools on long reads. With respect to computational performance, SeqScreen-Nano exhibited lower run time and reduced RAM usage compared to MetaMaps and MEGAN-LR, while being slower yet more memory efficient than Centrifuge. Notably, the spiked ZymoBIOMICS Microbial community DNA standard results indicate that the reference inference module using a breadth of coverage approach outperforms taxonomic classifiers that often have lower confidence results upon decreasing the abundance threshold for sample-wide analysis. Since estimating abundance thresholds for real metagenomic data is often difficult, reference-guided approaches represent a promising avenue to identify low-abundance pathogens in complex samples^38^. Apart from simulated and synthetic datasets, we also analyzed a real metagenomic dataset from a healthy FMT donor. Using our reference inference approach, we were able to identify several genera and species that are indicative of a healthy gut microbiome. Apart from *antibiotic resistance*, we also observed a trace amount of ORFs associated with *cytotoxicity* and *virulence activity*. Further research is needed to understand the interactive effects of different FunSoCs in the gut microbiome community towards clinical screening of FMT samples. Improved FMT screening is critical for protecting patient health as hard-to-characterize pathogenic microbes may be present within screened and approved FMT samples^39^.

Despite these promising results, SeqScreen-Nano is not without its limitations. While our experimental results highlighted that SeqScreen-Nano performs better in detecting the presence of individual pathogens within a given sample, it yielded lower precision and recall at read-level classifications. To partially account for this limitation, a larger set of possible taxonomic ids are transferred to the SeqScreen-Nano reference inference module from the ORF assignments, rather than the narrow set of single taxonomic ids from each read. We leave as future work additional refinements and improvements to read-level classification improvements, including integration with MetaMaps for mapping to nucleotide databases and exploring the use of outlier detection on bit scores^40^, as well as exploring adding a normalized bit-score based majority voting and weighted minimum set cover scheme. Moreover, previous studies have noted inherent challenges in using a gene catalog database (like UniRef100) for metagenomic analysis due to possible transitive clustering errors as well as multi-species clusters resulting in incorrect taxonomic assignments^41^. While we have tried to alleviate outlier assignments within gene clusters using our minimum set cover approach accounting for all taxids within the cluster, further improvements to the algorithm are still needed to increase accuracy.

In this work, we explore the challenging task of identifying previously unseen pathogens through a simulated experiment. While this may be considered as biologically unrealistic, often novel pathogens (in humans) involve the presence of specific genes that have evolved from near-neighbor pathogens to cross species barriers under a stringent evolutionary selection mechanism^42–44^. One such important mechanism is Horizontal Gene Transfer^45^. In our simulated experiments, we observed that simple gene (or sequence) transfer events from a pathogen to a near-neighbor can confound pathogen identification using taxonomic classifiers. We feel this is an important experimental observation and one that underscores the need for taxonomy-oblivious approaches like SeqScreen-Nano. Typically, standard k-mer based or phylogenetic placement based approaches tend to approach pathogen identification based on how the majority of reads, or the majority of the genome, are classified. While reasonable in many settings, these approaches might not always reflect “pathogenic potential” and virulence factors within the genome. SeqScreen-Nano identifies the presence of pathogens through FunSoCs that are based on key functions in biological pathways specific to pathogens. However, it is important to note that the FunSoC framework itself is limited to only identifying genes that are homologous to those present in the UniRef database, so novel genes that exhibit very distant homology (less than 50% aa similarity) to those contained in UniRef may be missed.

## CONCLUSION

We have highlighted the strengths and weaknesses of existing approaches for in-field pathogen detection, and have described them in detail. The SeqScreen FunSoC framework meets this challenge by enabling the detection of pathogenic genes in potentially novel pathogens. SeqScreen-Nano presents a novel paradigm for gene-based identification and labeling of pathogenic markers from long-read metagenomic data. It provides an efficient way to identify ORFs across the length of the read and assign functional and pathogenic labels to each ORF. Further, SeqScreen-Nano also offers a reference inference module based on ORF annotations to retrieve the most likely genomes present in the sample. This represents the first long-read bioinformatics capability to agnostically annotate ORF functions and determine the most likely microbial reference genomes present in a mixed community. Overall, the coupling of technological advances in the space of portable long-read sequencing and computational approaches for biosurveillance hold great promise to held prevent and mitigate future pandemics. In summary, SeqScreen-Nano presents a resource-efficient and accurate computational platform for rapid in-field detection of previously unseen pathogens.

## ACKNOWLEDGMENTS

The authors would like to thank Nicolette Keplinger, Matthew Scholz, Leslie Parke, and the broader Signature Science team for helping with the technical execution and administrative management of this project. The authors also express their appreciation to all of the early end users who provided constructive feedback for this software, including Dr. Cory Bernhards and his team at the U.S. Army CCDC Chemical Biological Center. All of the co-authors were either fully or partially supported by the Fun GCAT program from the Office of the Director of National Intelligence (ODNI), Intelligence Advanced Research Projects Activity (IARPA), via the Army Research Office (ARO) under Federal Award No. W911NF-17-2-0089. The views and conclusions contained herein are those of the authors and should not be interpreted as necessarily representing the official policies or endorsements, either expressed or implied, of the ODNI, IARPA, ARO, or the US Government. M.N. was partially supported by a training fellowship from the Gulf Coast Consortia, on the NLM Training Program in Biomedical Informatics & Data Science (T15LM007093). This work was supported in part by the Big-Data Private-Cloud Research Cyberinfrastructure MRI-award funded by NSF under grant CNS-1338099 and by Rice Universitys Center for Research Computing (CRC). T.T. was also supported in part by the National Institute of Allergy and Infectious Diseases (Grant# P01-AI152999) and by National Science Foundation grant EF-2126387.

## AUTHOR CONTRIBUTIONS

K.T and T.T managed and supervised the SeqScreen-Nano project. A.B implemented the methods and performed the experimental analyses. Y.L developed the reference inference module and assisted with the experimental evaluation. A.B., K.T., and T.T wrote the manuscript.M.N contributed to the streaming capability of the tool. B.H contributed to the development and benchmarking. A.K led the laboratory effort to sequence the Zymo gut microbiome community, tested the streaming capability and reviewed the FunSoC framework. D.L assisted with sequencing the ZymoBIOMICS Gut Microbiome Standard. G.G developed the FunSoC framework and reviewed results during tool development. All authors edited, read, and approved the manuscript.

